# Is it what’s inside that matters? A conserved microbiome in woody tissues of *Pinus radiata*

**DOI:** 10.1101/2023.08.10.552887

**Authors:** Kaitlyn Daley, Yeganeh Eslami, Charlotte Armstrong, Kathryn Wigley, Steven A. Wakelin

**Affiliations:** Scion, Private Bag 3020, Rotorua 3046, New Zealand; Global Plant Research Ltd, Mount Maunganui, Bay of Plenty, New Zealand; Scion, P.O. Box 29237, Riccarton, Christchurch 8440, New Zealand

**Keywords:** *Wood microbiome*, *Pinus radiata*, *bark*, *cambium*, *pith*, *carbon*

## Abstract

Understanding the interaction of endophytic microbiomes and their tree hosts may provide insights into wood formation and quality. Given the role of wood in carbon and nutrient cycling, this will provide valuable insights for forest growth and carbon cycling globally. Furthermore, the management of these interactions may add new value to wood-and fibre-based forest products. We assessed the microbiome of outer and inner bark, cambium tissue, year 2-8 wood increments, and the pith of 11 *Pinus radiata* trees, a widely planted, model conifer species. Diverse prokaryotic and fungal microbiomes were present in all trees, with communities structured by tissue type (p<0.001). Inner and outer bark tissues had high richness and the most distinct communities. Microbiome richness was lowest in year 2 through to year 8 wood, and the communities in these samples had similar composition. Prokaryote communities were dominated by Alpha-Beta-, and Gamma-proteobacteria, Actinobacteria, Firmicutes (Clostridia and Bacilli). Within fungal communities, Sordariomycetes comprised over 90% of the taxa present. Microbiomes of cambial and pith tissues were distinct to those niches. Overall, we provide further support that the wood of conifers is host to distinct microbiome communities. Microbiomes in these niches are profoundly placed to impact tree physiology, health, and fitness, through to ecosystem function and global carbon cycles.

## Introduction

Microbes and plants exist in a symbiotic partnership where the fitness of each is shared (Hawkes and Connor 2017; Sánchez-Cañizares et al. 2017). Increasing our understanding of these relationships is not only important for the plant but extends to the ecosystem it inhabits and the functions and services these ecosystems support (Brockerhoff et al. 2017). Studies in plant systems have demonstrated that microbes are a component of most plant tissues and that factors impacting the microbiome structure, function, or interaction with plant tissues, influence host physiology, and fitness (Bulgarelli et al. 2013; Stone et al. 2018; Vandenkoornhuyse et al. 2015; Warren 2022). These associations can be exploited to increase the quality and/or quantity of plant-based products and their sustainable production (Sivakumar et al. 2020), and delivery of a wide range of other ecosystem services spanning carbon storage, biodiversity support, water regulation and so forth.

The majority of plant-microbiome research has focused on the leaf and root compartments, the phyllosphere and rhizosphere, respectively. Understanding these niches is important for plant production and ecosystem productivity. These are, of course, the main tissues where the exchange of gasses, nutrients, and water occurs, and can be subject to disease which impacts overall plant production. Similarly, research on fruit and seed microbiomes has been important given potential for microbiomes to improve quality, alter traits, or increase production and security of supply of the food system (Hyde et al. 2019; Sivakumar et al. 2020). However, microbiome associations with woody plant stem and how these influence tissue physiology and plant fitness, are yet to be widely explored (Koskella 2020).

Approximately 550 gigatons of carbon is held within the Earth’s biosphere (i.e. across all life, in all habitats). Of this, an estimated ∼450 Gt is held in plants, with ∼300 Gt of this estimated to be held in tree stems (Bar-On et al. 2018). Thus, tree stems – *and principally woody tissues* – are estimated to hold in excess of half the total biomass carbon present on Earth. This is an unimaginably vast organic biome, a massive volume of habitat for microbes. Thus, not only is woody tissue the most neglected habitat of phytobiome research, but we posit that this is also one of the most understudied yet vitally important microbiomes globally *per se*.

The flow of carbon and energy into and through ecosystems, particularly forests, is a key driver of ecosystem functions (Odum et al. 1962). On this basis, influences on processes such as those regulating carbon flow with the biosphere or alter the chemical structure or composition of carbon pools will have profound impact on ecosystem-level processes (e.g. Smith et al. 2000).

These influences scale to impact Earth system processes. The extent of forests as terrestrial biomes globally (38% of global habitable land is forested; Ritchie and Roster 2021) and the size of the carbon pool held in woody-stem tissues, means that alteration of carbon exchange between woody biomass in the biosphere and atmosphere impacts climate change. Similarly, the flow of carbon into, within, and from this pool is central to the earth systems process, driving energy flow, nutrient cycling, supporting biodiversity, and ecosystem productivity and resilience. Factors such as chemical composition (e.g. degree of lignification or secondary metabolite deposition) that protect the wood from degradation and increase ecosystem residence time, have far-reaching consequences. The alteration of polyphenolic compounds can, as an example, extend the preservation of wood for hundreds of years (Jansson et al. 2010) dramatically altering the exchange of energy and resources within the biosphere and exchange with the soil and atmosphere pools. Yet unless forests are harvested for timber or fibre, the importance of wood is often overlooked. Despite this, the importance of wood formation and decay remains. Wood growth, for example, may have higher sensitivity to environmental parameters than photosynthesis, such that portioning of carbohydrates to non-structural v. structural pools transitions under stress. The incorporation of such processes into global vegetation models is needed as they underpin, again, processes that scale to Earth-systems level (Friend et al. 2019).

Wood is not a single or uniform tissue type but refers to a wide range of structural carbon tissues typified by lignified cell walls with wide ratios of carbon to other nutrients. In tree stems, woody tissues can include bark, the outmost protective layer of the stem, sitting adjacent to the vascular cambium. Bark provides a barrier of protection against fire, insect and pathogens, mechanical damage (e.g. Boland and Woodward 2021; Lawes et al. 2011), and other factors to the fragile, actively-growing outer layer of the stem; the vascular cambium. The exterior of the bark is also in direct contact with the environment and, as such, exposed to conditions other woody stem tissues are largely shielded from. The vascular cambium comprises highly active undifferentiated cells that divide to form phloem and xylem, the latter of which form the main woody part of the tree stem (Ye and Zhong 2015). The vascular network itself can constitute transport corridors for microorganisms to disseminate within the plant, connecting above and below ground biomes (Bahram et al. 2021; Yadeta and Thomma 2013). It provides sources of labile substrate (photosynthate) that can support microbial growth. Any microbial growth in the cambial zone may have profound impacts through the plant; this is where cell division, differentiation and activity occurs, a source and sink of hormones compounds, and a vascular network that provides a signalling network to root, leaves and tissues through the entire plant (Wang 2020). Microbial activity in this tissue is likely the production of hormones or other signalling molecules which is likely to directly impact processes related to wood formation and quality.

Further into the stem is the main, structural wood, primarily composed of ‘dead’ secondary xylem cells but also contain and abundance of living cells such as parenchyma, resin canals, and active epithelial cells (Schmitt et al. 2021). Finally, the centre of tree stems often contains pith, comprised largely of parenchyma tissue. Processes affecting the amount of pith material, and relations between pith and formation of reaction wood, can significantly impact downstream utilisation of wood, particularly log/timber product grades and value (Timell 1986).

Before we can start to investigate how microbes effect woody-tissue and how they can be manipulated to increase the productivity and other wood properties/processes, the presence of a microbiome within the different woody tissue needs to be confirmed. Therefore, the goal of this work was to determine (1) if DNA evidence for prokaryotic and/or fungal microbiome community could be detected within the woody tissue of a single *Pinus radiata* specimen with amplicon sequencing of genomic DNA, and (2) if any microbiome is present, is there any consistency (reproducibility) in community composition among trees indicating that wood-type habitats are selective.

## Materials and Methods

### Preliminary Sampling: Is there an inner sanctum?

Initial sampling focussed on determining if a potential microbiome is present in the ‘inner sanctum’ of *P. radiata* woody stem tissue. If present, it was considered that the microbes may derive from the root-soil system and then distributed vertically up the tree (e.g. via transport in the vascular networks). We hypothesised that there would be greater microbial richness lower in the tree, which could diminish with tree height. Therefore, collection of samples was also structured to test for influence of height.

Collection was conducted on a seven-year-old *P. radiata* tree, approximately 14 metres in height, with typical growth form and no apparent disease. The individual was growing in a small stand of *P. radiata* adjacent to the Whakarewarewa Redwood forest, Rotorua, New Zealand. Wood materials were collected along 50 cm height increments from the base of the tree up to 5 m, and then at 1 m steps further up. All instruments were sterilised with 70% ethanol between each sampling.

### Main collection: The microbiome at breast height (MBH; 1.37 m)

The second (main) collection of samples was collected from 11 clonally propagated trees (clone 15) in Kaingaroa forest (∼N571599, ∼E1892933). The trees were all in visibly good health and were growing within a uniformly managed single-age (12 years old) class stand. Based on findings for the previous samples (i.e. the inner sanctum data), collections of material for microbiome analysis were only made at breast height, or approximately 1.37 m; a standard heigh used for mensuration of tree trunks and therein allometry in forestry science (Magarik 2021). Trees were sampled while standing (i.e. not felled). Samples were collected from the south facing surface of each tree using sterile instruments as described above.

### Tissue processing, DNA extraction and quantification

Bark tissue was sampled using a wood window hammer. The material was then separated into the outer bark and inner bark surfaces. A wood corer was used to collect other wood samples from the exposed trunk surface (Fig. 1). This included the ‘cambium’, covering the true cambial meristem layer as well the broader cambium zone encompassing recently differentiated phloem and xylem cells. The wood core was sectioned and every second-years growth rings, up until 8 years, collected (e.g. Fig. 1). Following this, the pith material was identified and excised.

**Figure 1:**
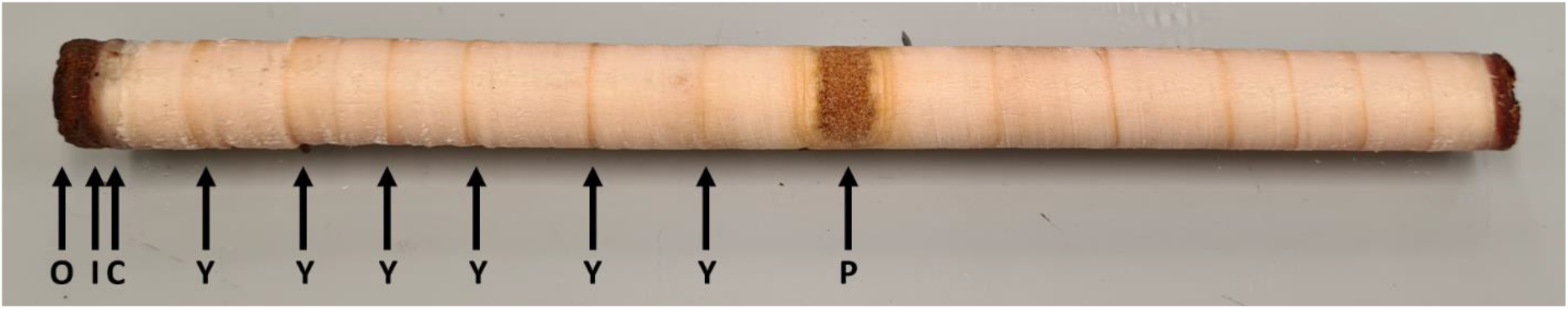
Example of a wood core extracted from *Pinus radiata*: (O) Outer bark section; (I) Inner bark section; (C) Cambium, white/translucent tissue; (Y) Yearly wood growth bands marked; (P) Pith at the centre of the stem. Note that ‘cambium’ includes true cambial cells as well as adjoining, recently differentiated phloem and xylem cells; our definition is operational and based on sampling practicalities.

Upon collection, bark samples were carefully wrapped in Parafilm and placed into a UV sterilised bag. Each core sample was individually inserted into an ethanol-sterilised plastic straw and covered in Parafilm. All samples were placed on ice and chilled for return to the laboratory. They were then stored at −20°C until processing for DNA extraction.

From each sample, 100 mg of tissues was taken and frozen in liquid nitrogen. Under sterile conditions, these were ground using a pestle in a mortar, then 50 mg transferred to 2 ml Plant Pro tissue disruption tubes (Qiagen Ltd, U.K.). DNA was extracted following manufacturers protocol, including the use of 50 μl of PS solution as recommended for difficult tissues.

All DNA was quantified using the Quant-iT PicoGreen fluorescence assay (Invitrogen Ltd, USA) on a Varioskan plate reader (Thermo Fischer Scientific, USA). Pith samples had low concentrations of DNA and were subsequently cleaned and concentrated into 35 µl using the Zymo Clean & Concentrate-25 kit (Zymo Research Corporation, Ca, USA). For all samples, working solutions were made at 2 ng DNA μl ^-1^ prior to PCR.

### Amplicon generation and sequencing

Sequencing of microbial communities followed methods described by the Earth Microbiome Programme (EMP; Thompson et al. 2017). In brief, for characterising the bacterial and archaeal communities, amplicon libraries based on the V4-V5 hypervariable regions of the 16S rRNA gene were created using primers 515F and 806R (Parada et al. 2016). For the fungal community, ITS1 rRNA libraries were generated using the primers ITS1f (Ihrmark et al. 2012) and ITS2 (White et al. 1990).

PCRs were conducted over 35 cycles and were based on TaKaRa Ex Taq Hot Start polymerase chemistry (Takara Bio, USA). For 16S rRNA gene amplification, conditions were as follows: 94°C for 3 minutes, followed by 35 cycles of 94°C for 40 seconds, 50°C for 60 seconds, 72°C for 90 seconds, and a final extension at 72°C for 10 minutes. Conditions were similar for fungal ITS1 rRNA amplification, except dissociation was for 30 seconds, primer annealing was conducted at 52°C for 30 seconds, and extension at 72°C for 30 seconds. After PCR clean-up, samples were pooled into equimolar solutions. Bacterial libraries were sequenced using 2 x 250 bp PE chemistry, and fungi 2 x 300 bp PE chemistry. Sequencing was conducted on Illumina MiSeq system at the Australian Genome Research Facility (AGRF).

### Bioinformatic processing of sequence reads

Sequences were filtered and trimmed using standard filtering parameters with DADA2 (Callahan et al. 2016). As part of the DADA2 pipeline, sequences were dereplicated, paired reads merged, and chimeras removed. Taxonomy was assigned to the sequences using the RDP trainset v18/release 11.5 for bacteria (Wang et al. 2007), and the UNITE database v8.3 for fungi (Nilsson et al. 2019). Sequences of plant-based origin (e.g. chloroplast) were removed from the datasets along with ASV’s unclassified at kingdom or phylum levels, or those present as singletons within a sample. Sequencing data for both the inner sanctum and MBH experiments is available in the NCBI Short Read Archive under bioproject accession number: PRJNA906136.

### Statistics

For alpha diversity assessment, the richness and evenness of ITS and 16S rRNA variants (at ASV level) in each sample were calculated using Margalef’s and Pielou’s indices, respectively. Values were assessed across tissue types using a one-way non-parametric ANOVA (Kruskal-Wallis), with t-testing among pairs of tissue types adjusted with Dunn’s-correction for multiple-comparisons (PRISM v9; GraphPad Software, CA, USA). Rarefaction curves were generated in R (R Cove team 2020) for individual samples to ensure that the depth of sequencing was sufficient to ensure collection of the rRNA variants present in each sample.

Taxa were aggregated from ASV to genera, family, and so forth to phylum. Resemblance matrices were created at each level using the Bray-Curtis distance method, selected due to its general applicability for ecological datasets and as it meets key guidelines for resemblance methods (Clarke et al. 2006). A 2^nd^ stage matrix was constructed and visualised using non-metric multidimensional scaling (nMDS; not shown). This enabled determination of the influence of taxonomic aggregation on distances among samples. From this, beta-diversity analyses were carried out at class level.

For the inner sanctum samples, variation in beta diversity was determined to assess if wood tissue type or sampling height was associated with variation in community composition. ASV counts were standardised across samples to minimise effects related to different total abundances, then square-root transformed to downweigh contribution of highly dominant taxa. Similarity among samples was calculated using the Bray-Curtis method (as before). PERMANOVA (Anderson 2001) was used to test for the effects of tissue type on community similarity (fixed term), with height of sampling represented as a covariate. Ordination via nMDS was used to visualise similarity in community type among samples. Where indicated, boot-strapping was used to calculate averages and variation (at 95%) for samples. Cluster analysis with SIMPROF testing (Clarke 1993; permutations = 999; alpha = 0.05) was used to explore if natural grouping of samples was present (i.e. not testing for statistical differences among *a priori* defined factors).

For the microbiome at breast height samples, all pre-treatment of data for beta diversity analysis was conducted as described above. However, PERMANOVA was used to test if microbiome communities in the wood tissue differed among the individual wood compartments across the 11 trees sampled (i.e. replication n = 11). Multivariate analyses were conducted using the PRIMER7/PERMANOVA+ package (PRIMER-E, Ltd., U.K.) using methods and approaches described previously (Anderson et al. 2008; Clarke and Gorley 2015).

## Results

### The microbiome of the Inner Sanctum

The complete panel of sequences were obtained from 76 tissue sections. After filtering and quality checks there were 1,548 and 2,997 unique ASVs present in the fungal and prokaryote datasets, respectively. In total, there were 3,786,878 sequences in the fungal dataset and 453,239 sequences in the prokaryote dataset. Rarefaction curves are given in the Supplementary Figures; Fig. S1A for prokaryote data, and Fig. S1B for fungal data.

Alpha diversity was assessed for sample collection height and tissue type. Summary plots are given in the Supplementary Figures S2A-S2D. ANOVA testing found no differences in prokaryote ASVs richness among tissue type (p=0.240), however evenness did vary (p<0.001). The summary pair-wise comparisons among the tissue types are given in Supplementary Table S1. For the fungal community, no differences in evenness were present among the samples (p=0.1). However, differences in fungal richness were evident (p<0.01). Pair-wise testing found these with inner bark samples had greater microbiome richness than samples of year 4 (p=0.02) or year 8 wood (p=0.02) samples.

Beta diversity summary treatment effects (PERMANOVA testing) are given in Table 1. Differences among wood tissues accounted for most variation in prokaryote ASVs community structure (p=0.001; √C.V.= 28.99; Table 1). These differences are most evident between the outer bark and internal wood tissues, and follow a progression from year 2 to year 8 wood (Fig. S3). However, the innermost wood sample, i.e. the pith, was unusual in having a bacterial community compositionally more similar to the outer bark (exposed tree surface) samples than those from internal woody samples (Fig. S3). No associations were evident for collection height nor interaction of height x tissue type and the variation on prokaryote community structure (Table 1)

**Table 1:**
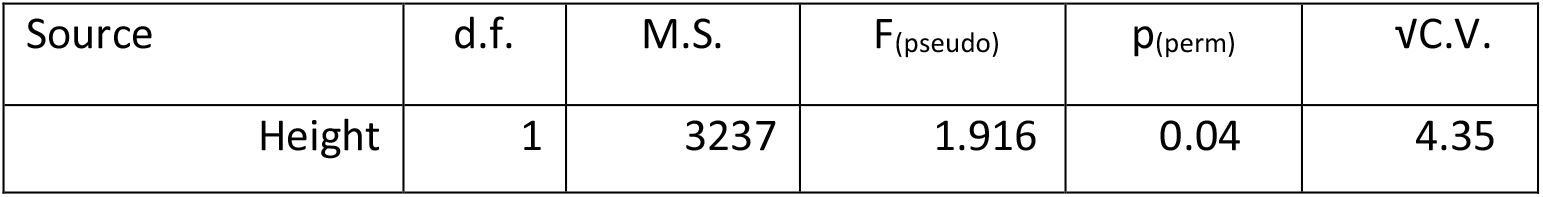
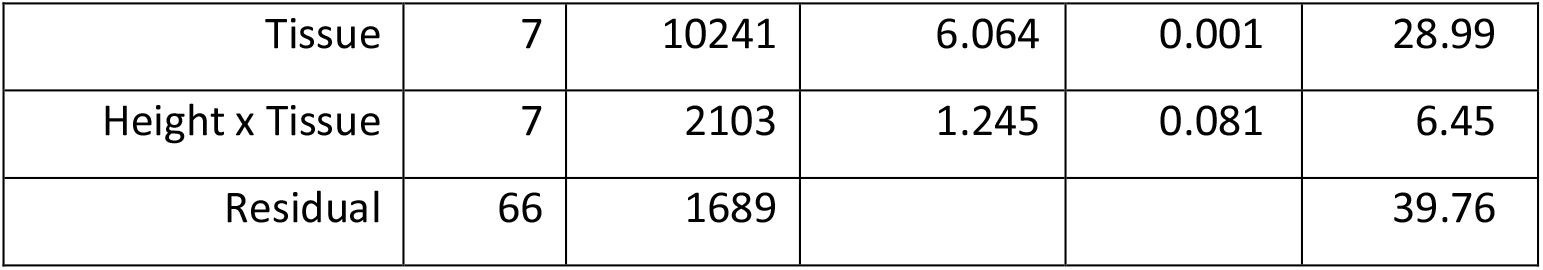
Summary PERMANOVA results for prokaryotes community structure in the Inner Sanctum.

The composition of fungal species also varied strongly among the different issue types (p=0.001, √C.V.= 21.99; Table 2). Changes in community composition are shown in Fig. S4. There is clear grouping of communities based on those from ‘bark tissues’ (outer bark, inner bark, cambium) when compared with those from the inner wood tissues (Fig. S4). For the fungal community, the pith community structure was more like the other internal woody tissues than the bark samples (contrasting with findings from bacteria; Fig. S3).

**Table 2:** Summary PERMANOVA results for the fungal community structure in the Inner Sanctum.

| Source | d.f. | M.S. | $F_{(pseudo)}$ | $p_{(perm)}$ | $\sqrt{C.V.}$ |
| --- | --- | --- | --- | --- | --- |
| Height | 1 | 1810 | 1.219 | 0.275 | 2.14 |
| Tissue | 7 | 5742 | 3.871 | 0.001 | 21.99 |
| Height x Tissue | 7 | 1940 | 1.308 | 0.098 | 7.26 |
| Residual | 55 | 15484 |  |  | 38.517 |

Vertical stratification of samples was not found to contribute to variation in fungal community beta diversity (p>0.2; Table 2). Similarly, no interaction was evident between collection height and wood tissue type (p>0.09; Table 2).

In brief, the detection of prokaryotes and fungal communities in all samples of wood collected, and following evidence of community-level structuring associated with tissue type, provided confidence that a microbiome naturally co-exists within the woody biomass of *P. radiata*. Consistency in community type among samples indicated this microbiome is likely to be – at least to some extent – influenced by deterministic assembly process. As such, results of this initial analysis led to further sampling with a focus on determining if the microbiome of the woody tissues showed stability when assessed multiple *P. radiata* trees.

### Part 2 – Microbiome at breast height

After filtering and quality checks there were 1,373 unique fungal ASVs and 7,100 prokaryote ASVs. The total abundance observed was 8,020,905 and 854,159 counts, respectively. The microbial communities were made up of 80 classes of prokaryotes and 30 classes of fungi. Rarefaction curves are given in the Supplementary Figure S5A for prokaryotic sequences, and Fig. S5B for fungal sequences.

The richness and evenness of both fungal and bacterial communities varied significantly among tissue types (Kruskal-Wallis ANOVA p<0.0001 for all). Summary plots of the data are given in Fig. 2A – 4D, and individual t-tests among tissue types in the Supplementary Table S2. Fungal and bacterial richness (Margalef’s d’) tended to be greatest in the outside tissues, and then decreased with distance of sampling into the stem (Fig 2B and 4D). Notably, however, richness was high in communities sampled from the pith compared with the year 2 to year 8 wood samples. While the richness trends for bacterial and fungal communities were similar, the patterns of evenness among tissue samples were not (Fig 2A and 2C). Fungal evenness was highest in the bark and cambium samples compared with those sampled from the wood. For bacteria, however, the wood samples (year 2-8) had higher evenness that samples collected from the bark (Fig 2C).

**Figure 2:**
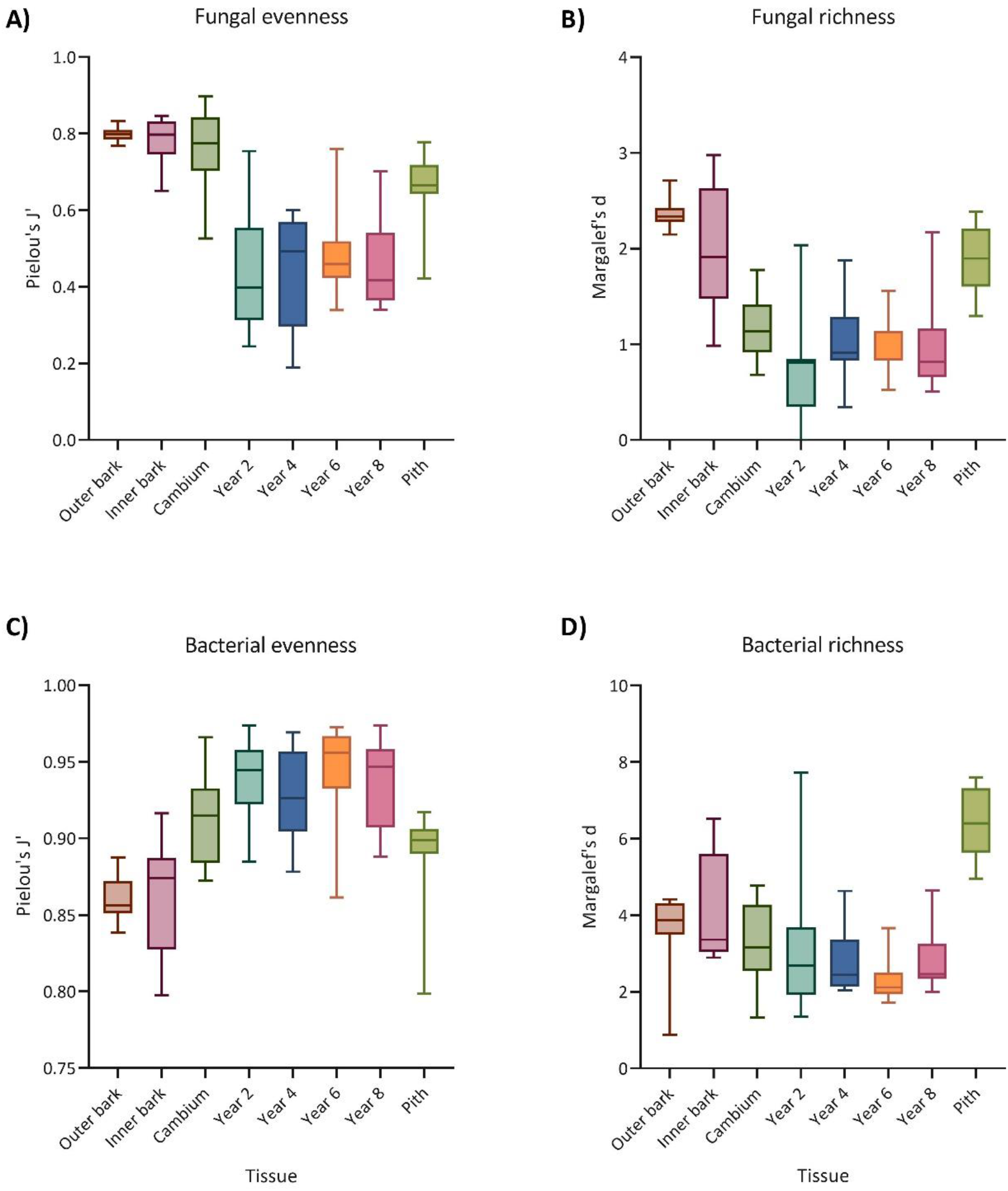
Evenness and richness of the microbiome at breast height. Each box gives median and range, with whiskers extending from minimum to maximum values for the 11 individual trees sampled. (A) Fungal ASVs evenness, (B) Fungal ASVs Richness, (C) Prokaryote ASVs Evenness, and (D) Prokaryote richness. Individual pair-wise tests among samples are not provided on these plots, but are provided in Summary Table S2.

### Prokaryotic beta diversity

Prokaryotic communities exhibited strong separation in composition among the main tissue type groups (p=0.001, √C.V.= 21.09; Fig. 3). The exception was the year 2 to year 8 woods; these grouped into an ‘inner wood’ type community (Fig. 3) with a generally shared microbiome. Clearer differentiation in community composition across the outer bark, inner bark and cambium tissues was present than observed for the inner sanctum samples (Fig. S3). This was likely due to a combination of the replication (i.e. statistical power) in the MBH samples, supported by sampling at a consistent height of 1.37 m. As for the bark and cambium tissues, the prokaryotic community in the pith sample was distinct from the other tissues (Fig. 3).

**Figure 3:**
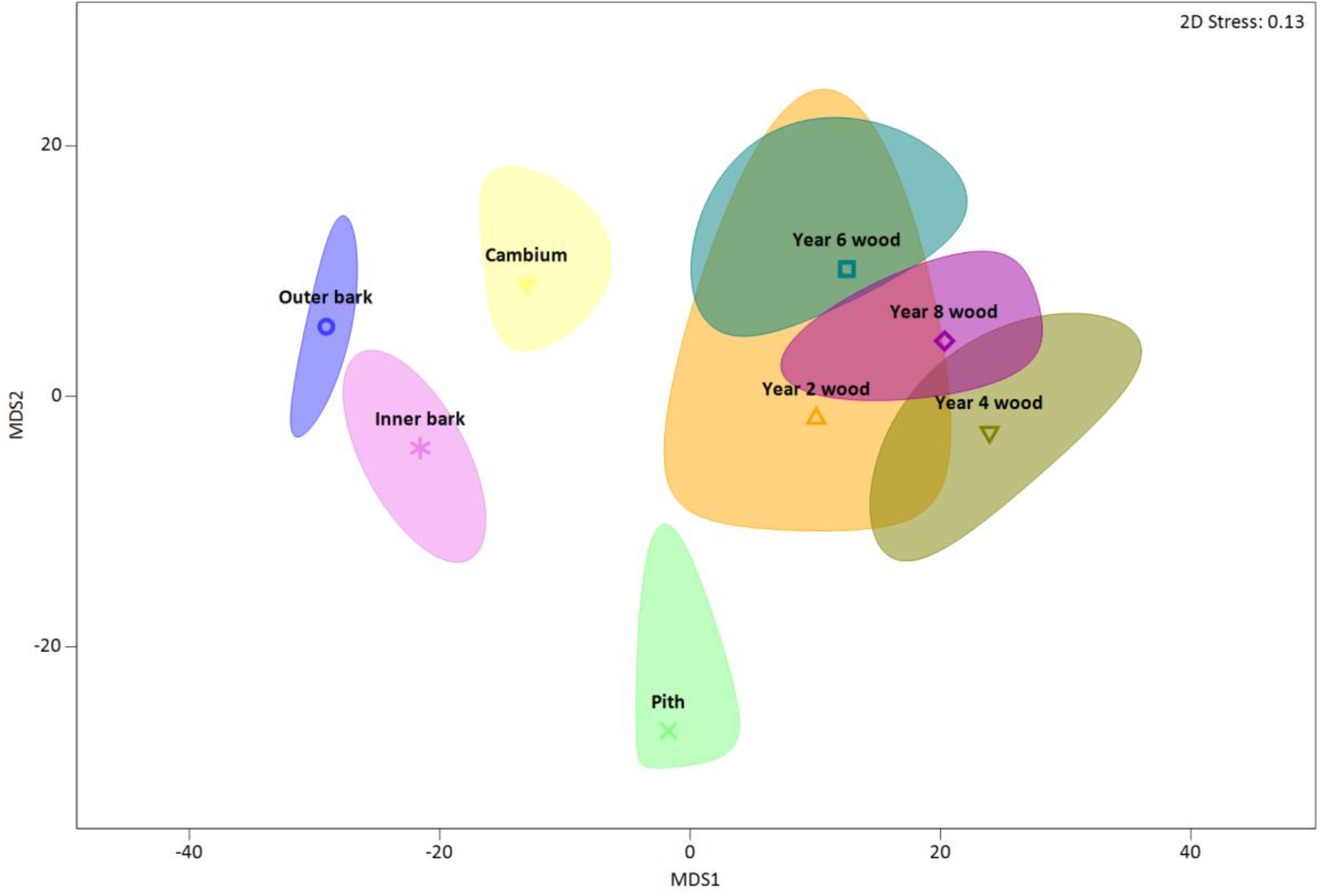
nMDS ordination showing similarity in prokaryotic community composition among tissue types for breast height collected samples. Points are boot-strapped averages for each tissue group, and shaded areas boot-strapped regions (95%). Results are at class level phylogeny, using square-root transformed abundance data, and differences calculated using the Bray-Curtis method

The relative abundances of the prokaryotic taxa found in the different wood tissues type are given in Fig. 4. Alphaproteobacteria and Actinobacteria comprised a main component of the *P. radiata* wood taxa, with the abundances of Actinobacteria increasing with samples collected further into the tree. The relative abundances of many taxa were related to tissue groups. For example, both Clostridia and Bacilli (both from the Firmicute phylum) were only found in significant abundances in the year 2 through to year 8 wood and pith. Similarly, Bacteroidetes were particularly abundant in the bark and cambium wood but were rare in other samples.

**Figure 4:**
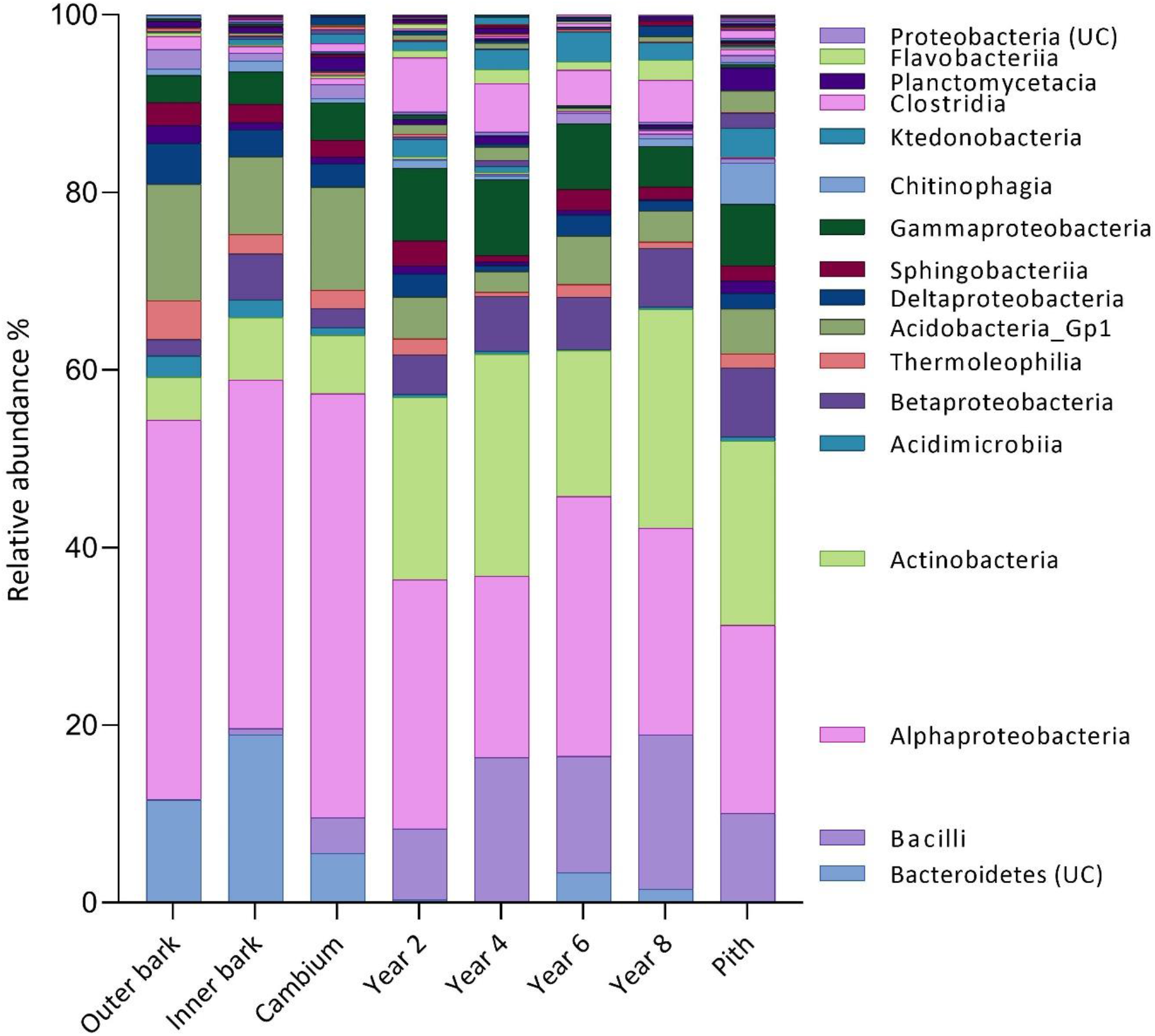
Relative abundances of prokaryotic classes averaged for each of the tissue types. Note the legend only shows major taxa. Labels with the suffix ‘UC’ indicate an unclassified group of that taxa.

### Fungal beta diversity

Variation in fungal communities among tissues broadly followed similar trends observed for prokaryotes; most wood tissue harboured a distinctive community type (p= 0.001, √C.V.= 27.51; Fig. 5), except for the year 2 to year 8 wood samples which overlapped. Furthermore, clear separation in community composition was evident among outer bark, inner bark, and cambium tissues (Fig 5). Overall, there is evidence for a step-wise transition in community type from the outer surface tissues though to the inner wood samples.

**Figure 5:**
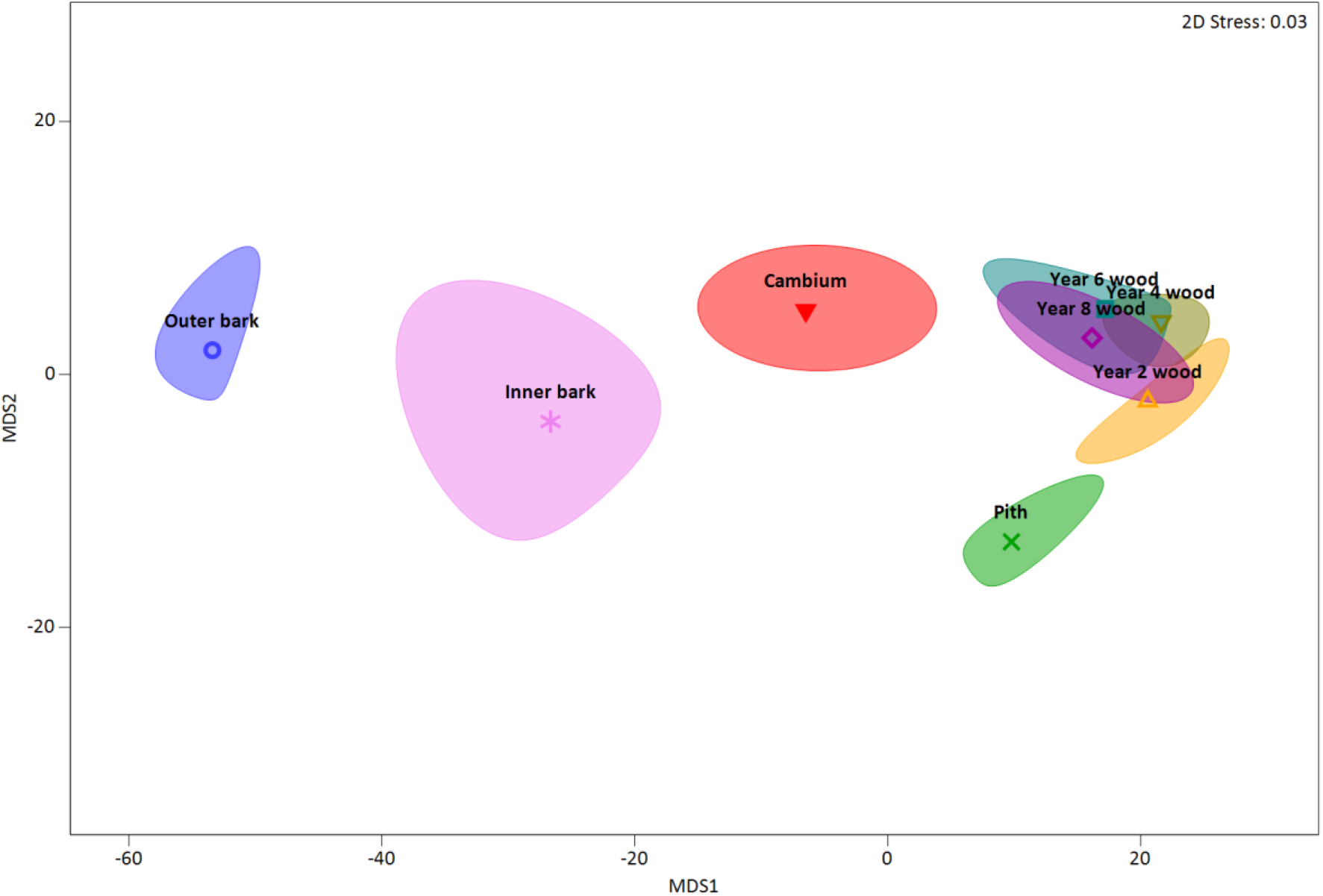
nMDS ordination showing similarity in fungal community composition among tissue types for breast height collected samples. Points are boot-strapped averages for each tissue group, and shaded areas boot-strapped regions (95%). Results are at class level phylogeny, using square-root transformed abundance data, and differences calculated using the Bray-Curtis method.

The relative abundances of fungal taxa in the wood tissues samples are given in Fig. 6. The outer tissue types held the most fungal taxa, and the abundances of the main groups – Sordariomycetes, Eurotiomycetes, an unclassified class of Ascomycota, Dothideomycetes, and Leotiomycetes – were more evenly spread across taxa present. With increased distance into the tree stem, the relative abundances of most fungal taxa declined while Sordariomycetes dominated (Fig. 6).

**Figure 6:**
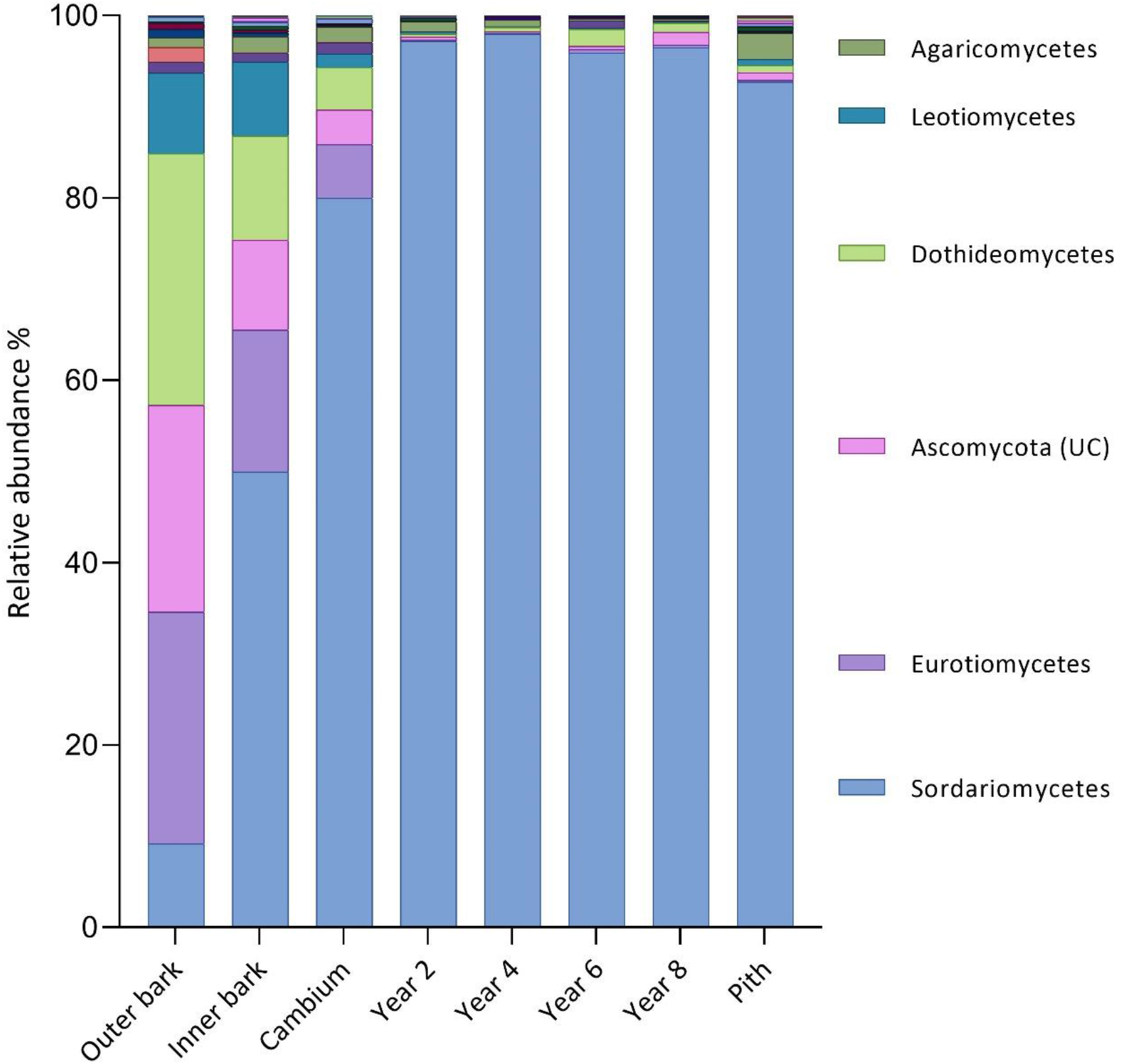
Relative abundances of fungal classes averaged for each of the tissue types. Minor taxa are not shown in the legend. Labels with the suffix ‘UC’ indicate an unclassified group of that taxa.

## Discussion

A rich community of microbes inhabit the wood tissues of *P. radiata*. Tissue samples collected from just above the ground level to more than 5 metres up the stem consistently hosted archaea, bacteria, and fungal communities. However, there was no apparent variation in community composition among these tissue habitats over the range of heights of sample collection. We hypothesised that if the source of wood microbiomes were from the soil/roots, then lower parts of the stem maybe richest in taxa which get ‘diluted’ in numbers and/or species with progression up the stem. This may be due, for example, to increased physical barriers (e.g. layers of cells) for the translocation of microorganisms, or chemical barriers and selection pressure by the plant, acting as a host filter. Although we found no strong evidence for this, it does not preclude the root from being a primary source for microbiome entry into the plant tissue. Rather, it means that (1) dissemination through the stem is subject to less filtering that we expected or, perhaps more likely, (2) that filtering had already occurred in the root tissue and subsequent dissemination of the microbes through the stem tissue isn’t restricted. Although transmission of microorganisms via living wood tissue occurs (Bahram et al. 2021; Yadeta and Thomma 2013), dissemination and niche filtering must still occur into the surrounding tissue, as each wood type harbours a distinct community.

Alternative avenues for establishment of the microbiomes in woody tissue include through direct environmental exposure. This can occur when lateral tissues, such as branches, emerge from initials (buds) beneath the bark, disrupting this natural barrier and exposing the inner stem surface. The integrity of bark as a barrier for the stem can, of course, be comprised due a number of biotic and abiotic events, from birds and insect damage, through to mechanical injury from wind and pruning. All present opportunities for exposure to and ingress of microbes from the environment. Whilst there are many potential routes for entry, we consider these are possible sources of microbiomes in this study. Although we sampled a visibly healthy stand of trees with no apparent bark damage, the stand of trees had been production pruned providing wounds and sites of potential entry for microbiomes. However, it could be expected that opportunistic entry into the tree by commensal microbes would result in limited potential to establish let alone disseminate through the unique habitat conditions of healthy wood tissue. In this case, if the wood endophyte microbiome was largely comprised of opportunistic taxa, we would expect to observe a reduced consistency in community assembly patterns across the microbiome among individual trees; i.e. community would be largely random to what opportunistically invades the tissue, not consistently structured in the species present as observed. Thus, we consider it likely that more selective mechanisms are occurring. The potential remains for entry from the root system and the distribution through the vascular tissue, particularly xylem (Bahram et al. 2021; Yadeta and Thomma 2013). It is also possible that generalised microbiomes are present in relatively undifferentiated stem tissue of seedlings from the nursery. When these are moved to the field and mature, filtering of microbes to the tissue habitats may occur. These all require further investigation and, for reasons discussed earlier, *P. radiata* is a useful model system for this.

We also urge care in overinterpretation of the finding that vertical height of tissue collection has no association with microbiome assembly. In this study, sampling was conducted from a single and relatively young tree. We would expect in taller and more mature trees that sampling height will be important. However, in the context of this study the initial ‘inner sanctum’ sampling was used to inform subsequent sampling of *P. radiata* trees similar-height and age. For this purpose, the finding was important.

The subsequent sampling across the 11 trees discovered taxa in wood tissue that were associated with mycorrhizal-associated fungi, soil-borne bacteria and archaea taxa (Paraburkholderia, Bradyrhizobium, Acidobacterium). The two classes of fungi that were only found in the pith tissues were Malasseziomycetes, and Wallemiomycetes. Both fungal taxa are found in a range of terrestrial environments including soil (Padamsee et al. 2012; Wang et al. 2014). A large abundance of different classes of Acidobacterium were spread throughout the prokaryote ASVs; this phylum thrives in low pH environments and, given the low expected pH of the wood (Hernández 2013), the presence of this phylum can be expected. Several prokaryote families found in the wood tissues include both aerobic and anaerobic genera. Families such as Prevotellaceae, Ruminococcacea, Lachnospiraceae are anaerobic prokaryotes, and were found everywhere with the exception of the outer bark samples. This finding itself demonstrates environmental-based filtering of community composition. Many of the groups of microbial taxa present within the woody tissue are broadly consistent with endophytes found various plant-microbiome systems. In particular, the presence of Alphaproteobacteria, Actinobacteria, Gammaproteobacteria, Firmicutes (Bacilli and Clostridia) among the prokaryotic community present (Hardiom et al. 2015), and the Sordariomycetes, Eurotiomyctes, and Dothidiomycetes in the fungal community (Hardiom et al. 2015).

Microbial communities in the inner and outer bark, cambium, and pith all showed strong similarity to each habitat (Fig. 3 and Fig. 5). However, samples from the year 2, 4, 6 and 8 wood increments overlapped, i.e. could not be separated on either fungal and prokaryotic community composition. Thus, five unique niches are present for *P. radiata*, and these are delimited by wood tissue type and not by distance into the tree. The presence of defined communities in each tissue indicates that, from a microbial perspective, they each comprise unique habitats. Although from a microbial perspective we do not know the factors on which the separation of the tissue groups into discrete habitats is based, it is reasonable to expect that differences in direct expose to environmental properties such as moisture, temperature, and UV exposure and, as we observed (see previously), aerobic v. anaerobic conditions, would disproportionately affect the outer wood samples (e.g. bark). Furthermore, while all wood samples maybe characterised by having wide ratios of C to other nutrients, chemical properties such as degree of lignification and extent and type of resins, tannins and other secondary metabolites may vary among the tissues. The importance of factors such as pH (Hernández 2013), seasonality and so on may also have a role in determining the composition and function of living wood microbiomes, yet these have only been the subject of a few studies (Beck et al. 2014; Krah et al. 2018; Ou et al. 2019). In particular, pH has been shown to be key factor associated with the microbiome composition in many ecosystems, including key plant compartments such as the phylloplane and rhizosplane (Dennis et al. 2009; Husson 2013; Gilbert and Renner 2021). The variation in pH of wood tissues may be an important factor associated with the observations of this study. It is known, for example, that pH of *P. radiata* sapwood can range from 3.83 to 5.7 depending just on the age of the tree (Hernández, 2013). It is unknown how variation in wood pH among different lineages of *P. radiata* or species of trees may affect this community, nor how historic selection of trees based on commercially desirable wood properties has impacted microbial communities present.

When in intimate association with and within plant tissues, microbial activity and interaction with the host tissue and metabolism can have profound influence on phenotype and fitness of the holobiont. Microbial production of metabolites associated with signalling compounds, hormones, toxins, and many other interactions can all modulate the plant metabolism (Hardoim et al. 2015). Furthermore, these do not need to be direct replication of host metabolites, but via chemical analogues that replicate or mimic plant signalling molecules, by impacting processes that dampen or amplify signalling cascades, altering transcriptional regulators and so forth. In short, the combination of potential influences from the microbiome against the opportunities for change in the host metabolisms presents an ‘interactomics hyperspace’ that is huge and effectively unexplored. However, it is also one of potentially immense value.

Agarwood provides a good example of where production of bioactive and secondary compounds by a microbiome can alter wood value. Agarwood occurs when wounds occurring in the stems of *Aquilaria* spp. are colonised by a specific microbiome that triggers stress-related secondary metabolite production and signalling (Naziz et al. 2019). Oleoresin is converted to wood resins, aromatic volatile compounds, and other secondary and biologically-active metabolites that stain and impregnates the wood tissue (Hyde et al. 2019). This resultant ‘Agarwood’ has uses from pharmaceutical/medical through to perfumes and is sought after globally. Indeed, it is one of the most expensive woods in the world and, due to illegal harvesting and smuggling, its trade is regulated under the Convention on International Trade in Endangered Species (CITES 2022). Overall, there is massive potential to utilise wood as a living bioreactor for production of microbial compounds. These span high-value secondary metabolites produced either by the microbes themselves, or the tree in response to the microbes, through to enzymes, alteration of wood for improved/altered feedstock properties for industrial purposes and so on (Hyde et al. 2019). How many interactomes are present in our trees, producing novel ‘forestceuticals’ is simply unknown.

Another opportunity is microbiome-based modification of wood to alter carbon residence time in ecosystems. For example, by reducing the rate of decay through infusion of wood with secondary metabolites, or by subtly altering the structural properties of wood (e.g. extent of lignification), this may increased residence time of carbon in the biosphere (Jansson et al. 2010). Given that more than half Earths biosphere carbon is estimated to be held in stem wood (Bar-On et al. 2018), this would have profound implications for global carbon cycling and climate change.

Future approaches, for example based on metatranscriptomics, are needed to determine the biomolecular activity of the different components of the wood microbiome over time, and how these influence the tree in changing environmental situations, across growth stages, during disease outbreaks, and other physiological conditions (e.g Nerva et al. 2022). In conjunction with understanding the fundamental ecology of the taxa present, how they are distributed in tissues, over time, and source and connection with the wider forest ecosystem, this may lead to new microbe-silvicultural opportunities to manage tree wood microbiome outcomes for benefit of forests, society and the environment.

## Acknowledgments

Dr. Yinon Bar-on, Weizmann Institute of Science, kindly conducted wood biomass carbon estimates based. Access to clone 15 *Pinus radiata* trees in Kaingaroa forest was kindly facilitated and supported by Timberlands Ltd. We thank the Scion field crew for support in tree processing and the testing of instruments for bark and wood collections. We appreciate the input of Dr Lloyd Donaldson, Scion, on comments and input on an earlier version of this manuscript.

