## Supplementary Table for "Is it what’s inside that matters? A conserved microbiome in woody tissues of *Pinus radiata*"

### Supplementary Tables

**Table S1.** Inner Sanctum Prokaryote (16S rRNA ASV's) richness (*d*) and evenness (J) summary statistics for t-tests between tissue types

| **Dunn's multiple comparisons test** | **Pielou (J') Evenness** | | | **Maraglefs (d) richness** | | |
|---|---|---|---|---|---|---|
| | Mean rank diff. | Summary | Adjusted P Value | Mean rank diff. | Summary | Adjusted P Value |
| **Outer vs. Inner** | 1.636 | ns | >0.9999 | -1.955 | ns | >0.9999 |
| **Outer vs. Cambium** | -2.273 | ns | >0.9999 | -9.455 | ns | >0.9999 |
| **Outer vs. Year 2** | -21.45 | ns | >0.9999 | 0.4545 | ns | >0.9999 |
| **Outer vs. Year 4** | -36.27 | * | 0.0137 | 4.818 | ns | >0.9999 |
| **Outer vs. Year 6** | -37.09 | * | 0.0102 | 17.00 | ns | >0.9999 |
| **Outer vs. Year 8** | -32.33 | ns | 0.2397 | 11.88 | ns | >0.9999 |
| **Outer vs. Pith** | -32.27 | ns | 0.0539 | 11.27 | ns | >0.9999 |
| **Inner vs. Cambium** | -3.909 | ns | >0.9999 | -7.500 | ns | >0.9999 |
| **Inner vs. Year 2** | -23.09 | ns | 0.6433 | 2.409 | ns | >0.9999 |
| **Inner vs. Year 4** | -37.91 | ** | 0.0053 | 6.773 | ns | >0.9999 |
| **Inner vs. Year 6** | -38.73 | ** | 0.0038 | 18.95 | ns | >0.9999 |
| **Inner vs. Year 8** | -33.97 | ns | 0.1385 | 13.83 | ns | >0.9999 |
| **Inner vs. Pith** | -33.91 | * | 0.0235 | 13.23 | ns | >0.9999 |
| **Cambium vs. Year 2** | -19.18 | ns | >0.9999 | 9.909 | ns | >0.9999 |
| **Cambium vs. Year 4** | -34.00 | * | 0.0228 | 14.27 | ns | >0.9999 |
| **Cambium vs. Year 6** | -34.82 | * | 0.0170 | 26.45 | ns | 0.3768 |
| **Cambium vs. Year 8** | -30.06 | ns | 0.3606 | 21.33 | ns | >0.9999 |
| **Cambium vs. Pith** | -30.00 | ns | 0.0878 | 20.73 | ns | >0.9999 |
| **Year 2 vs. Year 4** | -14.82 | ns | >0.9999 | 4.364 | ns | >0.9999 |
| **Year 2 vs. Year 6** | -15.64 | ns | >0.9999 | 16.55 | ns | >0.9999 |
| **Year 2 vs. Year 8** | -10.88 | ns | >0.9999 | 11.42 | ns | >0.9999 |
| **Year 2 vs. Pith** | -10.82 | ns | >0.9999 | 10.82 | ns | >0.9999 |
| **Year 4 vs. Year 6** | -0.8182 | ns | >0.9999 | 12.18 | ns | >0.9999 |
| **Year 4 vs. Year 8** | 3.939 | ns | >0.9999 | 7.061 | ns | >0.9999 |
| **Year 4 vs. Pith** | 4.000 | ns | >0.9999 | 6.455 | ns | >0.9999 |
| **Year 6 vs. Year 8** | 4.758 | ns | >0.9999 | -5.121 | ns | >0.9999 |
| **Year 6 vs. Pith** | 4.818 | ns | >0.9999 | -5.727 | ns | >0.9999 |
| **Year 8 vs. Pith** | 0.06061 | ns | >0.9999 | -0.6061 | ns | >0.9999 |

**Table S2.** Inner Sanctum Fungal (ITS rRNA ASV's) richness (*d*) and evenness (J) summary statistics for t-tests between tissue types

| **Dunn's multiple comparisons test** | **Pielou (J') Evenness** | | | **Maraglefs (d) richness** | | |
|---|---|---|---|---|---|---|
| | Mean rank diff. | Summary | Adjusted P Value | Mean rank diff. | Summary | Adjusted P Value |
| **Outer vs. Inner** | 4.614 | ns | >0.9999 | -7.314 | ns | >0.9999 |
| **Outer vs. Cambium** | -13.56 | ns | >0.9999 | -3.714 | ns | >0.9999 |
| **Outer vs. Year 2** | -14.83 | ns | >0.9999 | 6.922 | ns | >0.9999 |
| **Outer vs. Year 4** | -22.59 | ns | 0.7387 | 24.09 | ns | 0.5008 |
| **Outer vs. Year 6** | -11.59 | ns | >0.9999 | 17.69 | ns | >0.9999 |
| **Outer vs. Year 8** | -17.29 | ns | >0.9999 | 33.54 | ns | 0.2669 |
| **Outer vs. Pith** | -15.54 | ns | >0.9999 | 18.16 | ns | >0.9999 |
| **Inner vs. Cambium** | -18.17 | ns | >0.9999 | 3.600 | ns | >0.9999 |
| **Inner vs. Year 2** | -19.45 | ns | 0.8698 | 14.24 | ns | >0.9999 |
| **Inner vs. Year 4** | -27.20 | ns | 0.0899 | 31.40 | * | 0.0187 |
| **Inner vs. Year 6** | -16.20 | ns | >0.9999 | 25.00 | ns | 0.1893 |
| **Inner vs. Year 8** | -21.90 | ns | >0.9999 | 40.85 | * | 0.0230 |
| **Inner vs. Pith** | -20.15 | ns | >0.9999 | 25.48 | ns | 0.2595 |
| **Cambium vs. Year 2** | -1.273 | ns | >0.9999 | 10.64 | ns | >0.9999 |
| **Cambium vs. Year 4** | -9.027 | ns | >0.9999 | 27.80 | ns | 0.0574 |
| **Cambium vs. Year 6** | 1.973 | ns | >0.9999 | 21.40 | ns | 0.4941 |
| **Cambium vs. Year 8** | -3.727 | ns | >0.9999 | 37.25 | ns | 0.0559 |
| **Cambium vs. Pith** | -1.977 | ns | >0.9999 | 21.88 | ns | 0.6315 |
| **Year 2 vs. Year 4** | -7.755 | ns | >0.9999 | 17.16 | ns | >0.9999 |
| **Year 2 vs. Year 6** | 3.245 | ns | >0.9999 | 10.76 | ns | >0.9999 |
| **Year 2 vs. Year 8** | -2.455 | ns | >0.9999 | 26.61 | ns | 0.7620 |
| **Year 2 vs. Pith** | -0.7045 | ns | >0.9999 | 11.24 | ns | >0.9999 |
| **Year 4 vs. Year 6** | 11.00 | ns | >0.9999 | -6.400 | ns | >0.9999 |
| **Year 4 vs. Year 8** | 5.300 | ns | >0.9999 | 9.450 | ns | >0.9999 |
| **Year 4 vs. Pith** | 7.050 | ns | >0.9999 | -5.925 | ns | >0.9999 |
| **Year 6 vs. Year 8** | -5.700 | ns | >0.9999 | 15.85 | ns | >0.9999 |
| **Year 6 vs. Pith** | -3.950 | ns | >0.9999 | 0.4750 | ns | >0.9999 |
| **Year 8 vs. Pith** | 1.750 | ns | >0.9999 | -15.38 | ns | >0.9999 |

**Table S3. Richness and Evenness statistical summary of the MBH Fungi ASVs.**

| **Dunn's multiple comparisons test** | **Pielou** (J') Evenness | | | **Maraglefs (d) richness** | | |
|---|---|---|---|---|---|---|
| | Mean rank diff. | Summary | Adjusted P Value | Mean rank diff. | Summary | Adjusted P Value |
| **Outer vs. Inner** | 3.518 | ns | >0.9999 | 13.49 | ns | >0.9999 |
| **Outer vs. Cambium** | 7.152 | ns | >0.9999 | 38.42 | * | 0.0149 |
| **Outer vs. Year 2** | 49.02 | *** | 0.0001 | 53.18 | **** | <0.0001 |
| **Outer vs. Year 4** | 47.91 | *** | 0.0001 | 44.92 | *** | 0.0004 |
| **Outer vs. Year 6** | 43.18 | *** | 0.0009 | 46.18 | *** | 0.0003 |
| **Outer vs. Year 8** | 45.91 | *** | 0.0003 | 50.91 | **** | <0.0001 |
| **Outer vs. Pith** | 25.64 | ns | 0.3839 | 13.99 | ns | >0.9999 |
| **Inner vs. Cambium** | 3.633 | ns | >0.9999 | 24.93 | ns | 0.7813 |
| **Inner vs. Year 2** | 45.50 | *** | 0.0008 | 39.69 | ** | 0.0065 |
| **Inner vs. Year 4** | 44.39 | *** | 0.0009 | 31.43 | ns | 0.0822 |
| **Inner vs. Year 6** | 39.66 | ** | 0.0055 | 32.69 | ns | 0.0682 |
| **Inner vs. Year 8** | 42.39 | ** | 0.0020 | 37.42 | * | 0.0146 |
| **Inner vs. Pith** | 22.12 | ns | >0.9999 | 0.5000 | ns | >0.9999 |
| **Cambium vs. Year 2** | 41.87 | ** | 0.0052 | 14.76 | ns | >0.9999 |
| **Cambium vs. Year 4** | 40.76 | ** | 0.0056 | 6.500 | ns | >0.9999 |
| **Cambium vs. Year 6** | 36.03 | * | 0.0284 | 7.758 | ns | >0.9999 |
| **Cambium vs. Year 8** | 38.76 | * | 0.0114 | 12.48 | ns | >0.9999 |
| **Cambium vs. Pith** | 18.48 | ns | >0.9999 | -24.43 | ns | 0.8735 |
| **Year 2 vs. Year 4** | -1.109 | ns | >0.9999 | -8.258 | ns | >0.9999 |
| **Year 2 vs. Year 6** | -5.836 | ns | >0.9999 | -7.000 | ns | >0.9999 |
| **Year 2 vs. Year 8** | -3.109 | ns | >0.9999 | -2.273 | ns | >0.9999 |
| **Year 2 vs. Pith** | -23.38 | ns | 0.7909 | -39.19 | ** | 0.0078 |
| **Year 4 vs. Year 6** | -4.727 | ns | >0.9999 | 1.258 | ns | >0.9999 |
| **Year 4 vs. Year 8** | -2.000 | ns | >0.9999 | 5.985 | ns | >0.9999 |
| **Year 4 vs. Pith** | -22.27 | ns | 0.9028 | -30.93 | ns | 0.0958 |
| **Year 6 vs. Year 8** | 2.727 | ns | >0.9999 | 4.727 | ns | >0.9999 |
| **Year 6 vs. Pith** | -17.55 | ns | >0.9999 | -32.19 | ns | 0.0794 |

| | | | | | | |
|---|---|---|---|---|---|---|
| **Year 8 vs. Pith** | -20.27 | ns | >0.9999 | -36.92 | * | 0.0173 |

**Table S4 Richness and Evenness statistical summary of the MHB Prokaryote ASVs.**

| **Dunn's multiple comparisons test** | **Pielou** (J') Evenness | | | **Maraglefs (d) richness** | | |
|---|---|---|---|---|---|---|
| | Mean rank diff. | Summary | Adjusted P Value | Mean rank diff. | Summary | Adjusted P Value |
| **Outer vs. Inner** | -3.909 | ns | >0.9999 | -2.636 | ns | >0.9999 |
| **Outer vs. Cambium** | -31.09 | ns | 0.1208 | 8.545 | ns | >0.9999 |
| **Outer vs. Year 2** | -47.27 | *** | 0.0004 | 18.73 | ns | >0.9999 |
| **Outer vs. Year 4** | -42.73 | ** | 0.0025 | 19.73 | ns | >0.9999 |
| **Outer vs. Year 6** | -54.45 | **** | <0.0001 | 34.64 | * | 0.0413 |
| **Outer vs. Year 8** | -46.55 | *** | 0.0005 | 22.36 | ns | >0.9999 |
| **Outer vs. Pith** | -20.91 | ns | >0.9999 | -26.09 | ns | 0.4652 |
| **Inner vs. Cambium** | -27.18 | ns | 0.3524 | 11.18 | ns | >0.9999 |
| **Inner vs. Year 2** | -43.36 | ** | 0.0019 | 21.36 | ns | >0.9999 |
| **Inner vs. Year 4** | -38.82 | * | 0.0102 | 22.36 | ns | >0.9999 |
| **Inner vs. Year 6** | -50.55 | **** | <0.0001 | 37.27 | * | 0.0174 |
| **Inner vs. Year 8** | -42.64 | ** | 0.0025 | 25.00 | ns | 0.6086 |
| **Inner vs. Pith** | -17.00 | ns | >0.9999 | -23.45 | ns | 0.8767 |
| **Cambium vs. Year 2** | -16.18 | ns | >0.9999 | 10.18 | ns | >0.9999 |
| **Cambium vs. Year 4** | -11.64 | ns | >0.9999 | 11.18 | ns | >0.9999 |
| **Cambium vs. Year 6** | -23.36 | ns | 0.8952 | 26.09 | ns | 0.4652 |
| **Cambium vs. Year 8** | -15.45 | ns | >0.9999 | 13.82 | ns | >0.9999 |
| **Cambium vs. Pith** | 10.18 | ns | >0.9999 | -34.64 | * | 0.0413 |
| **Year 2 vs. Year 4** | 4.545 | ns | >0.9999 | 1.000 | ns | >0.9999 |
| **Year 2 vs. Year 6** | -7.182 | ns | >0.9999 | 15.91 | ns | >0.9999 |
| **Year 2 vs. Year 8** | 0.7273 | ns | >0.9999 | 3.636 | ns | >0.9999 |
| **Year 2 vs. Pith** | 26.36 | ns | 0.4344 | -44.82 | ** | 0.0011 |
| **Year 4 vs. Year 6** | -11.73 | ns | >0.9999 | 14.91 | ns | >0.9999 |
| **Year 4 vs. Year 8** | -3.818 | ns | >0.9999 | 2.636 | ns | >0.9999 |
| **Year 4 vs. Pith** | 21.82 | ns | >0.9999 | -45.82 | *** | 0.0007 |
| **Year 6 vs. Year 8** | 7.909 | ns | >0.9999 | -12.27 | ns | >0.9999 |
| **Year 6 vs. Pith** | 33.55 | ns | 0.0581 | -60.73 | **** | <0.0001 |
| **Year 8 vs. Pith** | 25.64 | ns | 0.5209 | -48.45 | *** | 0.0002 |

**Table S5. PERMANOVA summary results for microbiome at breast height analysis.**

| Source | d.f. | M.S. | F(pseudo) | P(perm) | √C.V. |
|---|---|---|---|---|---|
| **Prokaryotes** | | | | | |
| Tissue | 7 | 5580 | 8.14 | 0.001 | 21.09 |
| Residuals | 80 | 685 | | | 26.18 |
| **Fungi** | | | | | |
| Tissue | 7 | 8256 | 37.39 | 0.001 | 27.51 |
| Residual | 84 | 2201 | | | 14.86 |
