## Supplementary Figure for "Is it what’s inside that matters? A conserved microbiome in woody tissues of *Pinus radiata*"

### Supplementary Figures

**Figure S1. Rarefaction Curve of the Inner Sanctum wood microbiome samples.** Rarefaction plots for prokaryotic 16S rRNA and fungal ITS amplicons. A) Rarefaction curves for the prokaryotic 16S rRNA amplicons B) Rarefaction curves for the fungal ITS2 amplicon.

A)

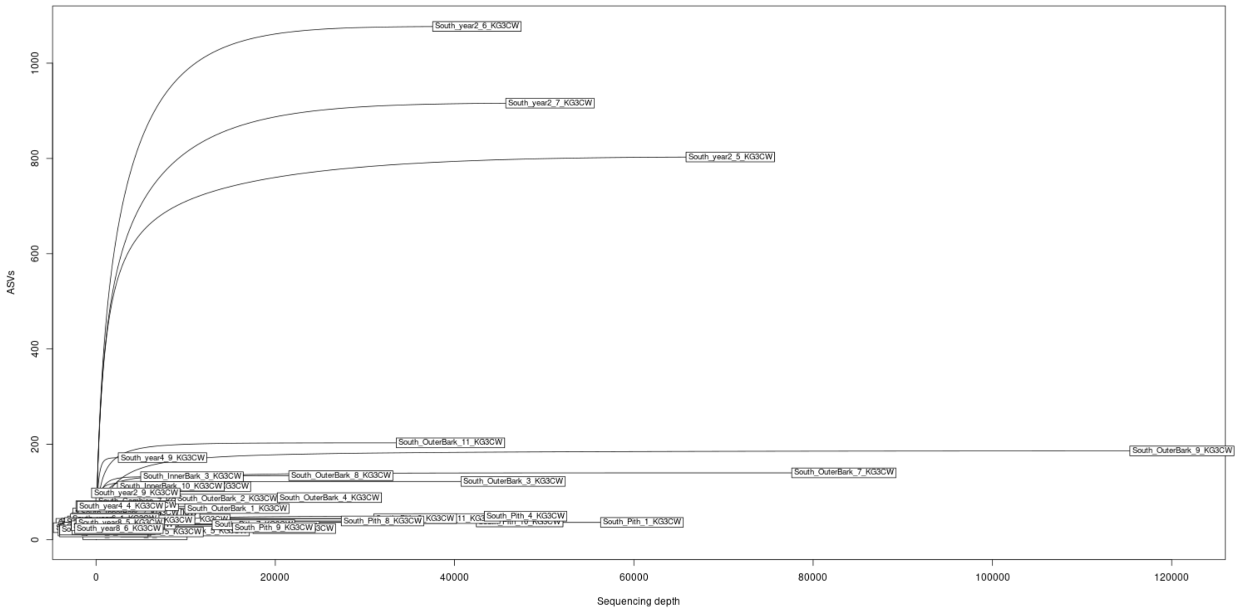


B)

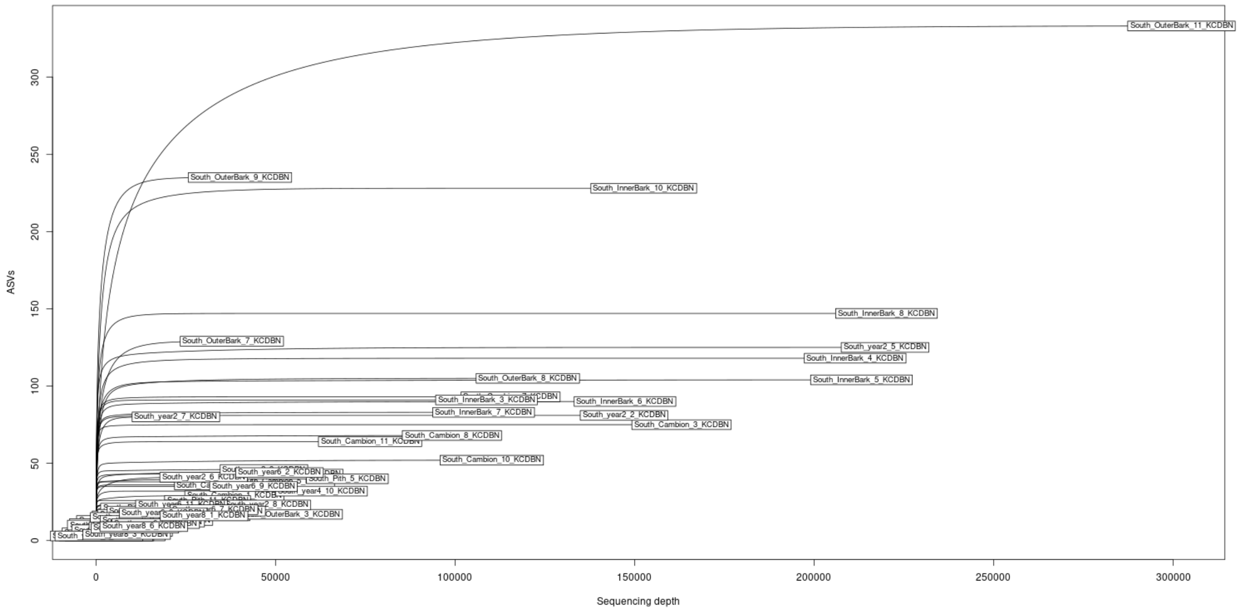

**Figure S2. Alpha diversity of the Inner Sanctum ASVs sectioned by wood tissue.** Pairwise comparison of the tissues showed only small significance (* p< 0.05). **A)** Fungal evenness (J'), **B)** Fungal richness (d), **C)** Prokaryote evenness (J'), and **D)** Prokaryote richness (d).

----*---- Prokaryotic evenness: Seven pairwise comparisons had p<0.05 and, due to difficulty plotting all of these, these are given in the Supplementary Table only.

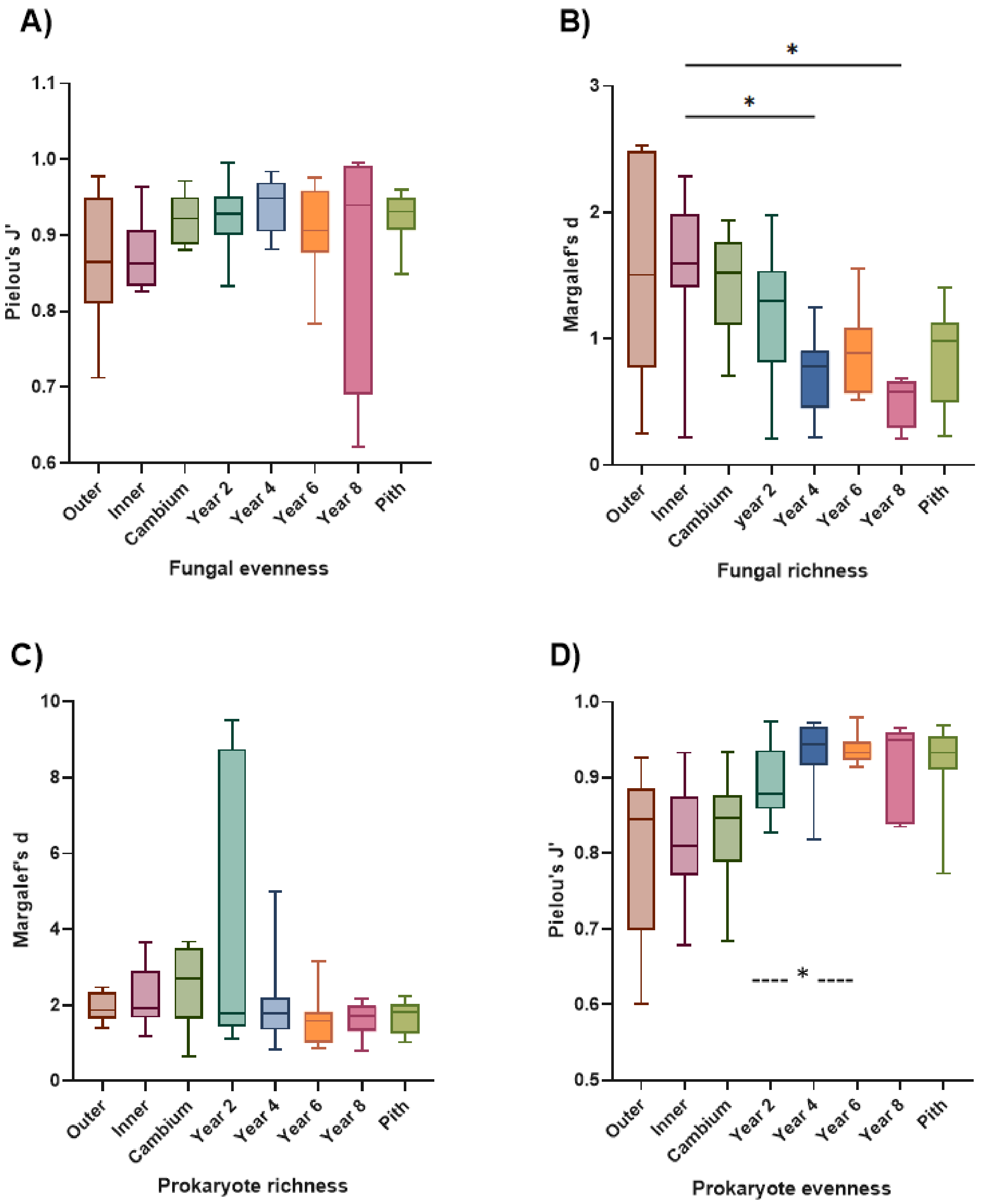

**Figure S3**: nMDS ordination showing similarity in prokaryotic community composition among tissue types. Points are boot-strapped averages for each tissue group, and shaded areas boot-strapped confidence regions (95%). Results are at class-level phylogeny, using square-root transformed abundance data, and differences calculated using the Bray-Curtis method.

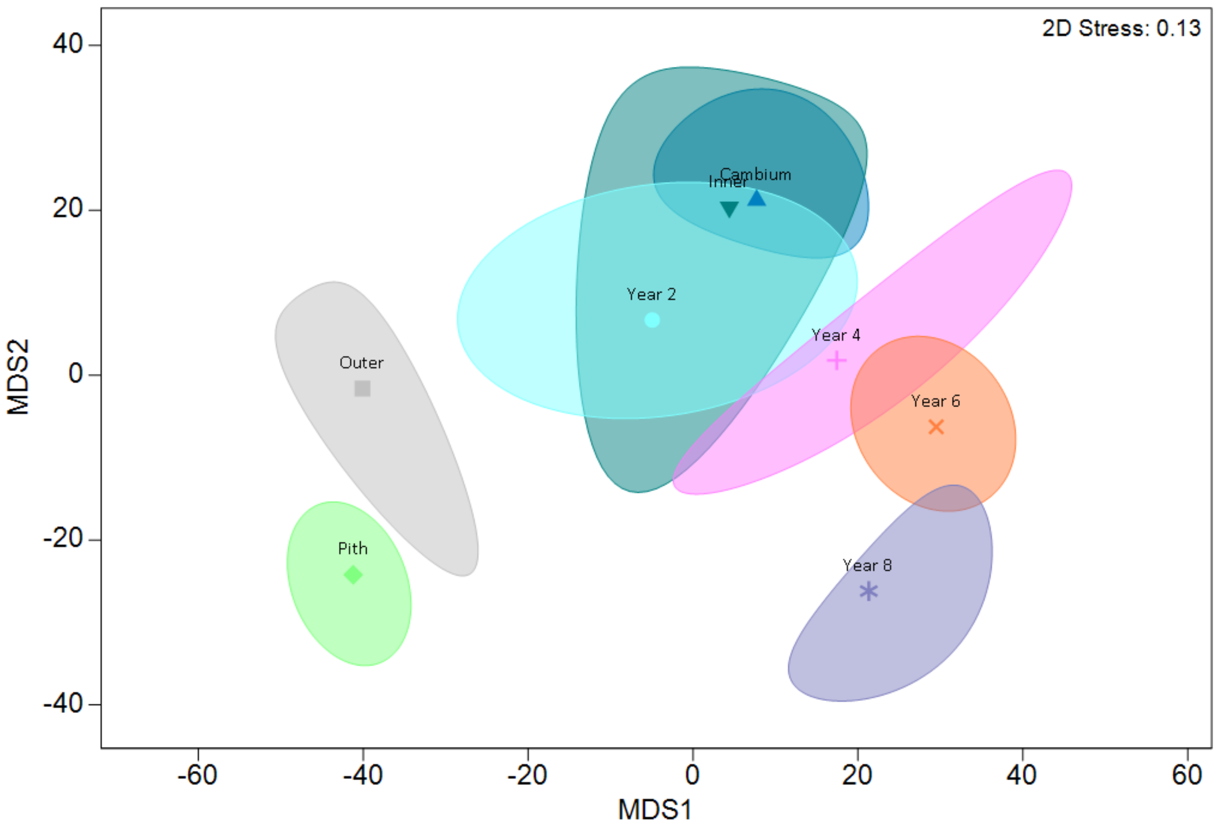

**Figure S4**: nMDS ordination showing similarity in fungal community composition among wood tissue types. Points are boot-strapped averages for each tissue group, and shaded areas boot-strapped confidence regions (95%). Results are at class-level phylogeny, using square-root transformed abundance data, and differences calculated using the Bray-Curtis method.

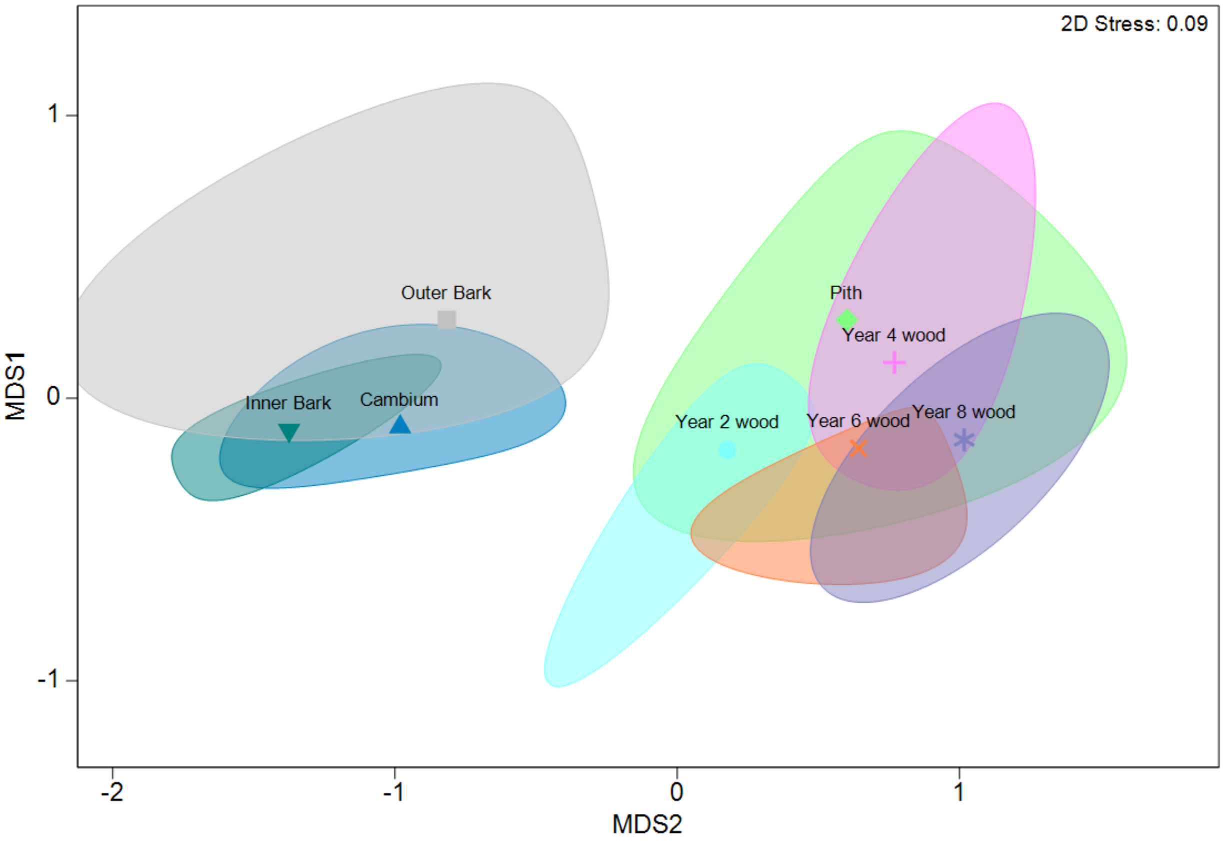

**Figure S5. Microbiome at breast height wood sample microbiome rarefaction curves.** Rarefaction plots for prokaryotic 16S rRNA and fungal ITS amplicons. A) Rarefaction curves for the prokaryotic 16S rRNA amplicons B) Rarefaction curves for the fungal ITS2 amplicon

A)

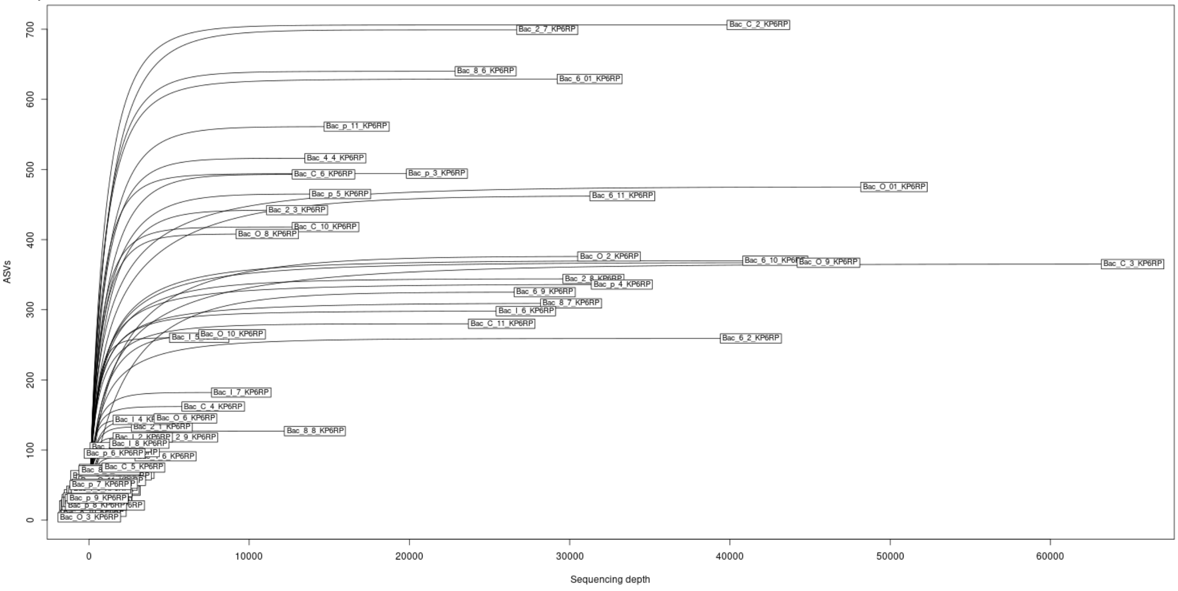


B)

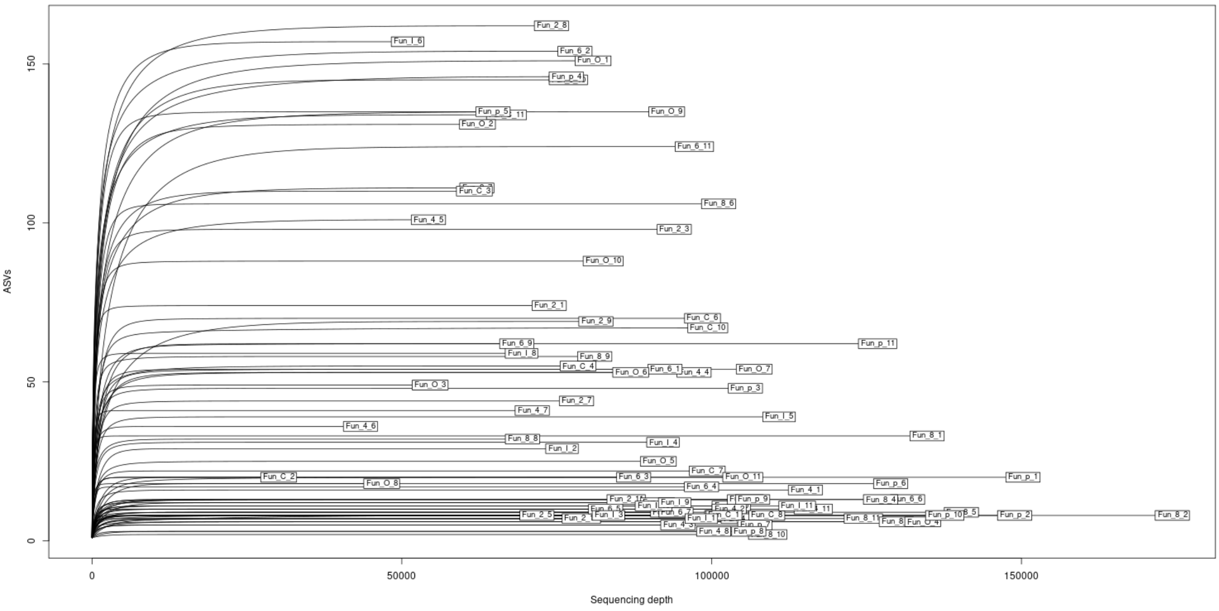
